# Evolutionary Genetics of Insecticide Resistance and the Effects of Chemical Rotation

**DOI:** 10.1101/072520

**Authors:** Dylan J. Siniard, Michael J. Wade, Douglas W. Drury

## Abstract

Repeated use of the same class of pesticides to control a target pest is a form of artificial selection that leads to pesticide resistance. We studied insecticide resistance and cross-resistance to five commercial insecticides in each of six populations of the red flour beetle, *Tribolium castaneum*. We estimated the dosage response curves for lethality in each parent population for each insecticide and found an 800-fold difference among populations in resistance to insecticides. As expected, a naïve laboratory population was among the most sensitive of populations to most insecticides. We then used inbred lines derived from five of these populations to estimate the heritability (*h*^2^) of resistance for each pesticide and the genetic correlation (*r*_*G*_) of resistance among pesticides in each population. These quantitative genetic parameters allow insight into the adaptive potential of populations to further evolve insecticide resistance. Lastly, we use our estimates of the genetic variance and covariance of resistance and stochastic simulations to evaluate the efficacy of “windowing” as an insecticide resistance management strategy, where the application of several insecticides is rotated on a periodic basis.

## Introduction

The adaptive response of pest arthropods to insecticide application is a striking example of rapid Darwinian evolution, occurring over a few generations in response to a strong man-made selection pressure. The ability of arthropods to evolve insecticide resistance has led in extreme cases to crop failures or resurgence of vector-borne diseases (Mallet 1989; Kranthi et al 2002). A better understanding of the phenotypic and genetic basis of the evolution of insecticide resistance and cross-resistance is necessary to slow resistance evolution and maintain control over pest insects.

Repeated use of the same class of pesticides to control a target pest is a form of artificial selection that leads to pesticide resistance. Worldwide, more than 500 species of insects, mites, and spiders have developed some level of pesticide resistance. The majority of currently commercialized synthetic insecticidal chemistries can be grouped into four modes of action: nerve and muscle function disruption, growth inhibition, respiration inhibition, and midgut disruption. Of the insecticides approved for use, the vast majority act on an organism’s neuro-musculature. The common physiological target of theses insecticides suggests that a pest’s response to treatment with one insecticide will be genetically correlated with its susceptibility to other functionally related insecticides. There is a large body of evolutionary genetic literature describing how to estimate such genetic correlations and how to use them to predict the correlated response of traits in populations, be it positive or negative, to bouts of selection (e.g., Lynch and Walsh 1998).

Here, we report our phenotypic and genetic estimates of resistance and cross-resistance to five commercial insecticides in each of six populations of the flour beetle, *Tribolium castaneum*. We use inbred lines derived from five of these populations to estimate two quantitative genetic parameters, the heritability (*h*^2^) and genetic correlation (*r_G_*) of resistance between pesticides in each population. These quantitative genetic parameters allow insight into the adaptive potential of populations to evolve insecticide resistance. Heritability determines how rapidly a population responds to a single selective agent, while genetic correlations determine how adaptation to one selective agent indirectly affects adaptation to other selective agents (Sih et al. 2004). In particular, positive genetic correlations imply that the evolution of resistance to one insecticide results in the evolution of cross-resistance to one or more other insecticides. Negative genetic correlations imply that the evolution of resistance to one insecticide results in an evolutionary increase in the sensitivity to one or more other insecticides. By estimating the genetic variance and covariance of resistance to insecticides with different modes of action across several populations, we are able to detect geographic variation in the capacity to evolve resistance. As we report, the data suggest that some populations have a past history of strong selection for resistance to some insecticides but not others. We use our estimates of the genetic variance and covariance of resistance to evaluate the efficacy of the insecticide resistance management strategy of “windowing,” wherein the application of several insecticides is rotated on a periodic basis to maintain pest-control efficacy and to delay the evolution of resistance.

Insecticide resistance tends to be polygenic when studied in laboratory selection experiments but monogenic when studied in pest insects isolated from the field (Roush and McKenzie 1987; ffrench-Constant et al. 2004). This finding has been attributed to the difference in the strength of selection imposed in the laboratory relative to that in the field. Because selection is generally weaker in laboratory evolution studies, they may favor resistance based on many genes, each of small effect, segregating within the experimental population. Field selection, in contrast, consists of applying much higher concentrations of insecticide and therefore stronger selection. As a result, the field response tends to be based on rare mutations of large effect at single genes (see Figure 1 in ffrench-Constant et al. 2004). Laboratory selection occasionally produces monogenic resistance, most often when genetic material is incorporated into the laboratory strain from field populations that have already had extensive exposure to the specific test insecticide or to an insecticide with cross resistance to the test insecticide. Although this dichotomy between monogenic and polygenic response may be an artifact of genetic interference (see below), we estimated the genetic variance and covariance of our populations for resistance at both the median lethal dose (LD_50_) and lethal dose, 90% (LD_90_) concentrations of all insecticides. This allowed us to compare the capacity for an evolutionary response at two different strengths of selection.

**Figure 1.**
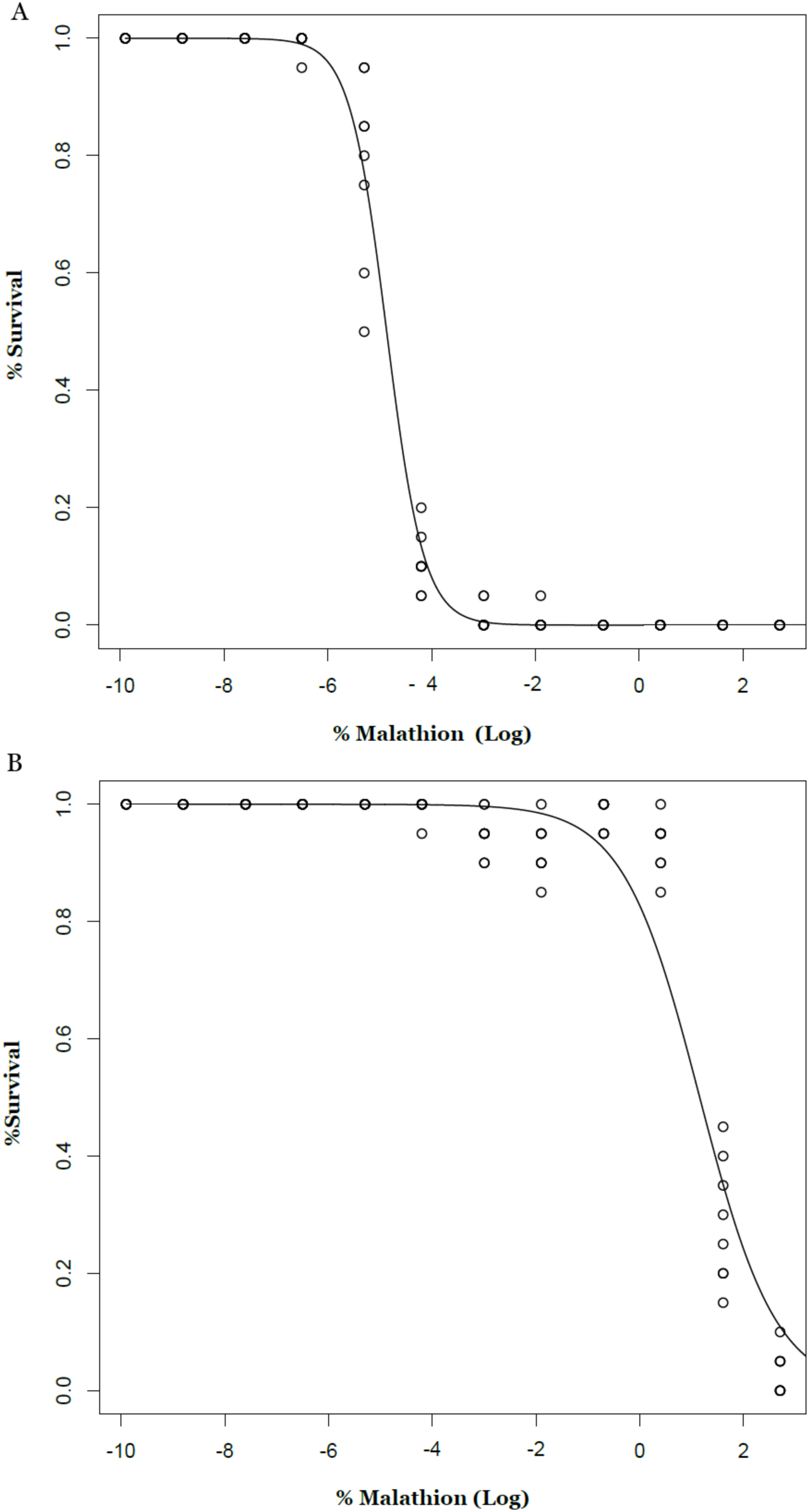
Estimation of the Malathion LD50 for the c-SM laboratory population (A) and Bhopal population (B). At the dosage of 0.0106 (SE: 0.0004) and 4.441 (SE: 0.166), there is 50% survivorship of c-SM and Bhopal adults respectively. A curve, as shown above, was generated for each population on each insecticide at 2 different time points, 18 and 65 hours. See supplementary material for all 60 curves.

In general, the initial response of a population to strong selection depends upon either the advent of new mutations or upon pre-existing alleles of major effect segregating within a target population (Olson-Manning et al. 2012). When dependent on new mutations, an adaptive response to strong selection is slowed by the waiting time necessary for those mutations to occur. Because the evolution of insecticide resistance has been so rapid, it is likely based on existing allelic variation, which in theory facilitates the most rapid adaptive response to strong selection (Olson-Manning et al. 2012). However, when two independently acting genes are subjected simultaneously to strong selection, ‘interference’ occurs between them, slowing the response to selection (Barton 1995). Interference occurs between simultaneously favored alleles at two loci because linked beneficial mutations arising in different individuals compete when the favored allele at one locus is found by chance in a linked genetic background with the non-favored allele at the other locus. Increasing the recombination rate between the two loci increases the likelihood that both favorable alleles will become fixed by selection. However, with strong selection even unlinked loci exhibit interference with one another (Barton 1994). Thus, when selection is strong, as it is for insecticide resistance in the field, interference occurs regardless of the chromosomal distribution of the selected alleles across a genome, because strong selection creates associations even between unlinked loci. Moreover, interference is asymmetric: genes of larger effect exert greater interference on genes of smaller effect. Thus, very strong selection may result in a selection response based on one gene of large effect even when other genes capable of contributing to the adaptive response are segregating in a population. In addition, interference is relatively more important to the response to strong selection in spatially structured populations, like agricultural fields, because local associations between loci created by selection are broken down more slowly because recombination is reduced by the deviations from random mating imposed by population structure.

We report our studies of resistance and cross-resistance to five commercial insecticides (Malathion, Sevin, Spinosad, Pyrethrin, and Imidacloprid) in the flour beetle, *T. castaneum*. In our studies, we used a global distribution of populations (South America, North America, Europe, India, and Africa) and a laboratory strain (c-SM), sequestered in the laboratory before 1940, i.e., before any of the tested commercial insecticides were developed. From all populations except that from Peru, we also derived a population of inbred lines by twelve generations of brother-sister mating. It is unlikely that flour beetles, which are common pests of stored products harvested for human consumption, have been targeted by direct application of any of the insecticides we studied. However, empty grain bins are routinely treated with insecticides, like methoxychlor, malathion or methoprene, prior to grain storage as a deterrent to flour beetle colonization. In addition, one of our experimental populations was collected in Bhopal, India, a few years after the explosion of a carbaryl insecticide manufacturing plant. In another population (Peru), a decades’ long mosquito abatement campaign, involving repeated application of pesticides, may have inadvertently affected *T. castaneum* (Griffing et al. 2013).

In this report, we first characterize the dosage response curves for lethality in each parent population for each insecticide. Notably, we discovered nearly an 800-fold difference among populations in resistance to some insecticides and, as expected, the naïve laboratory population was among the most sensitive of populations to most insecticides. Second, we report our estimates of the heritability (*h*^2^) and genetic correlation (*r_G_*) of resistance and cross-resistance, respectively, within populations (e.g., Jackson et al. 2007; Tabashnik et al. 2009). These estimates were obtained by testing the insecticide sensitivity of inbred lines around the LD_50_ and LD_90_ of the parent populations from which the lines were derived by inbreeding. Lastly, we use our estimates of the genetic variance and covariance of resistance to evaluate the efficacy of “windowing” as an insecticide resistance management strategy, where the application of several insecticides is rotated on a periodic basis.

## Materials and Methods

### Commercial Insecticides

We tested the response of populations of *T. castaneum* to commercial formulations of five insecticides encompassing five different classes of compounds. We purchased insecticides directly from distributors (see Table 1 for specifications of each formulation). All formulations act to disrupt neural transmission with the primary physiological targets being enzymes within the cholinergic pathway, except for pyrethrin which targets voltage-gated ion channels (ffrench-Constant et al 2004). Acetylcholine is a neurotransmitter that functions to send impulses between cells at muscular junctions. Nerve transmission is terminated by the cleaving of acetylcholine by acetylcholinesterase (AChE). Organophosphate and carbamate insecticides bind AChE irreversibly, preventing the termination of nerve transmission and resulting in death by hyper excitation and paralysis (ffrench-Constant et al 2004). Newer insecticides, such as spinosad and imidacloprid, act as antagonists of the nicotinic acetylcholine receptor (nACR), also resulting in death by hyper excitation and paralysis. The acetylcholine receptor is composed of 11 subunits, which are bound differently by each insecticide (Perry et al 2011). Newer insecticide formulations achieve their specificity by binding one or more of these subunits. Overall, these chemicals have rapid and robust insecticidal effects prior to the evolution of resistance.

**Table 1.**
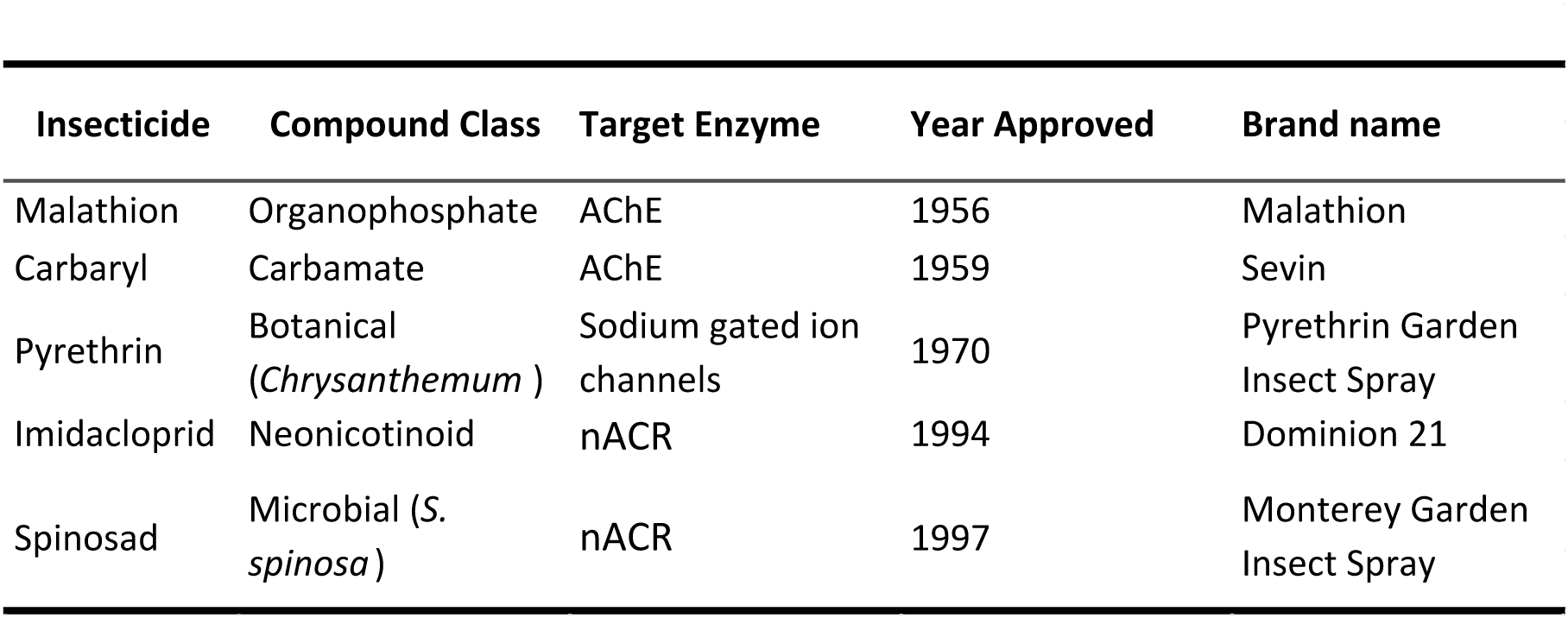
Insecticides used in this study.

### Species Description

*Tribolium castaneum*, a major global pest of grain and stored products, is a model laboratory organism used to develop and test important ecological and evolutionary concepts (e.g., Wade 2016). This species is believed to colonize grain stores through anthropogenic movement because its own dispersal abilities are limited (Drury et al. 2016; but see Ridley 2011). The combination of near obligate commensalism and limited dispersal results in a globally distributed species composed of a multitude of genetically differentiated populations with an average, pairwise, genomic *F_ST_* of 0.180 (Demuth and Wade, 2007 a and b; Drury et al., 2009).

We used beetles from six populations: five populations have a wide global distribution (South America, North America, Europe, India, and Africa) and one is a long-held laboratory strain (c-SM). The origin of the outbred laboratory population, c-SM, is described in Wade (1977). The other four populations were collected from various types of stored products over the last several decades (Table 2). All wild caught collections originated from more than 50 non-virgin adults. Subsequent stocks were maintained at a population size of > 500 individuals on a standard medium (20:1, flour: brewer’s yeast, by weight) and in 24 h darkness.

**Table 2.** *Tribolium castaneum* wild Population Information.

| Population | Collection Date | Collection Location |
| --- | --- | --- |
| Bhopal, India | 1989 | Flour Granary |
| Chicago, Illinois (c-SM) | 1938 | Laboratory Strain |
| Dar es Salaam, Tanzania (Des) | 1989 | Market |
| Jerez, Spain | 1998 | Livestock Feed |
| Purdue, Indiana, USA | 2008 | Livestock Feed |
| Lima, Peru | 1996 | Market |

### Inter-population Covariation Bioassays

We assessed the resistance of adult beetles from each population in Table 2 to each of the five insecticides in Table 1. We tested the lethality of Malathion and carbaryl at 12 concentrations of each insecticide: 0.00005%, 0.00015%, 0.00050%, 0.001500%, 0.005000%, 0.01500%, 0.05000%, 0.1500%, 0.50000%, 1.500%, 5.000% and 15.000%. Since lethality was essentially 0.00 at the five lowest concentrations, we eliminated the four lowest concentrations from our tests of pyrethrin, imidacloprid and Spinosad. Since commercial concentrations of Spinosad and pyrethrin were 0.50% and 1.0%, respectively, we did not investigate the higher concentrations for either insecticide.

We administered an insecticide by pipetting 2 mL of a solution at the desired concentration onto 400 mg of whole grain flour in a small, 20 mL weigh boat and allowed the medium to dry overnight into a chip. For each population and each dilution, we presented a chip to each of eight replicate groups of 20 adults (6 populations × 8.2 dilutions on average × 6 treatments (5 insecticides + control) × 8 replicates × 20 adults = 44,928 exposed beetles). Control media consisted of one chip with 2 mL of distilled water, which we considered a 0.00 ppm dilution in our analysis (see Analysis Section below). We recorded the number of dead adults in each replicate 18 and 65 hours (139 hours for pyrethrin) after the adult beetles were introduced to a treated chip. From the mortality data of these dilution series, we estimated the LD_50_ and LD_90_ for each population to each insecticide (see Analysis Section below).

### Intra-population Covariation Bioassays

From all populations but Peru, we created dozens of inbred lines by imposing twelve generations of brother-sister mating. We assessed the levels of genetic variation (heritability) and covariation (genetic correlations) of insecticide resistance by treating adult beetles from 12 inbred lines from each population with each insecticide at the LD_50_ concentration of the corresponding, outbred parent population. As above, we pipetted 2 mL of insecticide solution onto 400 mg of whole grain flour in a weigh boat and allowed the medium to dry overnight into a chip. We presented one chip to each of eight replicate groups of 20 adults (5 populations × 12 inbred lines × 6 treatments (5 insecticides + control) × 8 replicates × 20 adults = 57,600 exposed beetles). Controls for each line consisted of one chip with 2 mL of distilled water, considered a 0.00 ppm dilution as above. For a set of inbred lines, we recorded the numbers of dead adults in each replicate at the same time point used to calculate the LD_50_ of their respective outbred parent population.

For the Purdue and Bhopal populations, we estimated the genetic variation (heritability) and covariation (genetic correlations) of insecticide resistance using the inbred lines as above but at the LD_90_ concentration of each outbred population. This study allowed us to compare the heritabilities and genetic correlations of the LD_50_ and LD_90_ environments for these two populations.

### Analysis

For each insecticide at each observation time, we used a 4 parameter logistic regression to find the LD_50_ or LD_90_ and its corresponding standard error for each population. The duration of exposure in subsequent analyses was determined from observing the highest concentration that killed all treated individuals and the lowest concentration that killed no individuals. This ensured the LD_50_ would fall between these concentrations.

### Statistical Methods: Variance Components and Heritabilities

We estimated heritability of resistance for each population from the inbred line means using standard quantitative genetic methods (Lynch and Walsh 1998). When a sample of inbred lines is large and chosen randomly with respect to the character to be analyzed, then differences among line means provide an unbiased estimate of the genetic differences among lines for the characters. Under the assumption that the inbred lines are homozygous at the contributing loci, strain differences must be related to additive genetic variance and not to dominance interactions within loci. The nature of the relationship of strain differences to additive genetic variance can be deduced from the discussion of Crow and Kimura (1970, p. 100) concerning the effects of inbreeding on character variance. The additive genetic variance among inbred lines [V_a_(i)] is related to the additive genetic variance in the randomly mating population [V_a_(r)] from which they were derived by the coefficient of inbreeding, f: V_a_(i) = V_a_(r)(1+f). Thus, for high levels of inbreeding (i.e., f ∼ 1), the additive genetic variance in the population from which the inbred lines were derived can be estimated as half the variance among the inbred line means. Narrow sense heritability (*h*^2^) is the fraction of the total variance due to additive genetic effects. We estimated the narrow sense heritability from the inbred line values as *h*^2^ = ½Var(Among Inbred Line Means)/[½Var(Among Lines)+ Var(Within Lines)]. The within-line component of variance is a repeated measure, estimated from multiple, near genetically identical, individuals for each trait.

We calculated the heritabilities from the observed among-line variance components estimated from a generalized linear model with a binomial fit, using restricted maximum likelihood (REML), followed by a delete-one-line jackknife. The jackknife has been shown to perform well for estimating variance components, heritabilities and genetic correlations (Knapp et al., 1989; Simons and Roff, 1994). In the delete-one jackknife, all observations from one line are deleted and a ‘pseudo-value’ of the desired statistic is calculated. This process is repeated, until a pseudo-value has been created for the set of inbred lines for each population. The estimate of the statistic and its standard error are calculated from the mean and standard error of the full set of pseudo-values (Roff, 2006).

### Statistical Methods: Genetic Correlations for Resistance

We used standard analysis of variance procedures to partition the variance of measured resistance to different insecticides within and between inbred lines. The same standard procedures will yield a partitioning of the covariance for two traits if a synthetic variable, the sum of values for two characters, is formed for each animal in every line and subjected to the same analysis. The component of variance among lines for this synthetic variable contains the component of variance among lines for each of the two characters singly plus twice their among-lines component of covariance. This component of covariance is the bivariate analogue of the component of variance among strains.

### Matrix Stability

The genetic variance-covariance matrix (the ‘g-matrix’) summarizes the inheritance of multiple, phenotypic traits. The stability of this summarizing parameter is important as it affects our ability to predict how the phenotypic traits evolve by selection. To determine the stability of the covariance matrices among our populations, we compared our matrixes by the method of random skewers (Cheverud and Marroig 2007; Calsbeek and Goodnight 2009; Pitchers et al. 2013). After a set of random selection vectors is applied to each matrix in a pair of covariance matrices, the vector correlation between the altered matrixes is measured. The strength of this correlation, compared to the correlation of the random vectors, is a measure of the similarity of the matrices. In addition to estimating the stability of g-matrices among populations, we also estimated the stability of the LD_50_ and LD_90_ matrixes within populations.

### Truncation selection

Response (*R*) to short-term selection is the function of heritability (*h*^2^) and the selection differential (*S*): *R* = *h*^2^·*S*. This relationship is often called the breeders’ equation. The between-generation change, (the response to selection) *R*, is the change in means between the population before selection and the population in the next generation. *S* is the within-generation change in the mean due to selection as *S* = *µ*\* − *µ* where *µ* is the population mean before selection and *µ\** the mean of the parents that reproduce (the population mean after selection). Thus, *S* is the difference between selected parents and the population as whole (within generation). In the case of pesticide application, *S* can be related back to the dose of insecticide applied. Assuming a normally distributed polygenic basis for resistance, susceptibility should range from 0-1 with a mean of 0.50. If the population is treated at its LD_50_ concentration, the mean before selection will be 0.50 and the phenotypic mean of the individuals remaining post exposure will be the mean of the normal distribution as if it was truncated at 0.50 and 1. Thus, we can calculate the phenotypic mean of the post exposure individuals using the truncated normal distribution. Let u be the phenotypic mean of the population, sigma be the calculated phenotypic standard deviation, a be the lower bound of the distribution, the truncation point, and let b be the upper bound (in this case 1). The *Z*-scores associated with the distribution are:

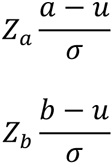

And the estimated phenotypic mean of the surviving individuals is then:

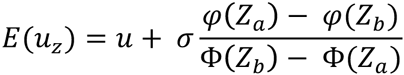

where ϕ is the probability density function of the standard normal distribution, and Φ is the cumulative distribution function of the standard normal distribution. Thus the selection differential *S* is equal to the difference between phenotypic mean of the selected population (E[*u_z_*])) and the unselected phenotypic mean (*u*), to give

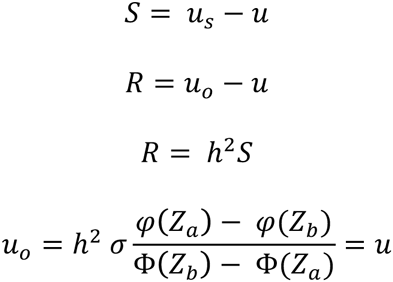

From this, we can estimate *u*_*o*_ the between generational change in LD_50_.

### Change in Correlated Characters

The extent to which traits are genetically correlated allows us to predict the genetic change in trait *Y* when truncation selection is applied a different but correlated targeted trait *X*. In this case, let *Xp* be the population phenotypic mean for lethality to a particular pesticide, and let *Yp* be the population mean for lethality to a second pesticide to which this population, this generation is not exposed. We calculate *Xo* using the equations above. To determine what effect the pesticide application will have on the population’s resistance to the unapplied pesticides we calculate the correlated change, where σ_*Ax*_ and σ_*Ay*_ are the additive genetic standard deviations of the applied and the untreated pesticides, respectively. And, *pG_xy_* is the additive genetic correlation of resistance phenotypes between the two chemicals:

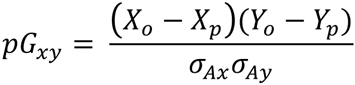

### Stochastic simulations

We ran evolutionary projections with the equations above and the observed quantitative genetic parameters estimated from our breeding experiments. The additive genetic variance, environmental variance and heritabilities were generated at random from a truncated normal distribution with a mean equal to the empirically estimated variance or heritability, a standard deviation equal to standard deviation of the jackknifed among-line pseudo-values of the inbred lines, and a truncation point at 0 to prevent the sampled variances from becoming negative. Phenotypic variances for the simulation were the sum of the sampled additive and environmental variance values. Phenotypic means for the first generation of simulations were the measured means from our population work. In generations beyond 1, the phenotypic mean from the previous generation (n-1) was used. For simulations in which only one insecticide is used, each generation, a new random draw from the distribution of possible V_A_, V_E_, *h*^2^ is used, the series of new means are followed for a total of five generations. To produce 95% predictive intervals around these estimates, the simulations are run 100,000 times and the bottom 2.50% and top 97.50% are used as the range of the 95% predictive interval.

We also ran simulations to estimate evolutionary trajectories when insecticides with known genetic correlations are rotated among generations. We used the same procedure as above, except two trait means were followed simultaneously. Each with its own V_A_, V_E_, and *h*^2^distributions, however, truncation selection is only applied to one trait each generation. The change in the alternative trait is estimated using genetic change in correlated characters equations. As with the single trait simulations, we use the genetic correlation and its error and the additive genetic variance in the alternative trait as the sampling distributions for these estimates.

## Results

### Variation among populations

We discovered a wide range of resistance profiles among our populations and among insecticides. Populations varied nearly 800-fold in susceptibility to Malathion with the Bhopal population (LD_50_: 4.441, SE +/− 0.166, n = 1920) and the Peru population (LD_50_: 5.576, SE +/− 0.168, n = 1920) exhibiting the highest resistance. The Jerez (LD_50_: 0.007, SE +/− 3.72 × 10^−4^, n = 1920) and the laboratory strain c-SM (LD_50_: 0.011, SE +/− 4.26 x 10^−4^, n = 1920) had the lowest resistance. The differences among populations in their resistance to Sevin showed a similar but less extreme pattern, with the Bhopal population (LD_50_: 1.417, SE +/− 0.088, n = 1540) and the Peru population (LD_50_: 2.633, SE +/− 0.139, n = 960) again on the high end of the resistance profiles and Jerez (LD_50_: 0.786, SE +/− 0.051, n = 1700) and the laboratory strain c-SM (LD_50_: 0.956, SE +/− 0.050, n = 1680) on the susceptible end.

For Imidacloprid, the most recently introduced of the insecticides tested, we observed intermediate variation in resistance among populations with a 5-fold difference between the highest and lowest resistance profiles. Again, we found that Bhopal (LD_50_: 0.945, SE +/− 0.063, n = 1280) and Peru (LD_50_: 0.898, SE +/− 0.049, n = 1280) were the most resistant populations but Tanzania (LD_50_: 0.161, SE +/− 0.011, n = 1280) and Purdue (LD_50_: 0.222, SE +/− 0.013, n = 1280) were the two most susceptible populations, more susceptible than the sheltered laboratory population c-SM (LD_50_: 0.395, SE +/− 0.022, n 1280).

All populations were highly sensitive to the certified organic insecticides and there was little variation among-populations in LD_50_. For pyrethrin, the LD_50_ concentration showed a 2.6-fold difference among populations with Peru (LD_50_: 0.039, SE +/− 0.002, n = 600) and c-SM (LD_50_: 0.033, SE +/− 0.002, n = 600) being the most resistant and Tanzania (LD_50_: 0.016, SE +/− 0.001, n = 600) and Jerez (LD_50_: 0.015, SE +/− 0.001, n = 600) being the most susceptible. Spinosad, a natural product of the soil bacterium *Saccharopolyspora spinosa* and relatively recently approved as an insecticide (DATE), exhibited the lowest variation in LD_50_ among populations with Peru (LD_50_: 0.149, SE +/− 0.009, n = 800) and Purdue (LD_50_: 0.096, SE +/− 0.006, n = 800) being the most resistance and Jerez (LD_50_: 0.069, SE +/− 0.004, n = 800) and c-SM (LD_50_: 0.070, SE +/− 0.006, n = 800) being the most susceptible.

### Genetic variation segregating within populations

We detected substantial variation among populations for resistance to all pesticides, the largest difference being the 796-fold difference in Malathion LD_50_ between the Bhopal and Jerez populations. This variability suggests that the different histories of exposure to insecticides have resulted in different levels of evolved insecticide resistance. We then turned to the question of the extent the observed phenotypic variation around the population mean estimates (represented above by their respective standard errors) was a result of genetic variation segregating among individuals within a population. To estimate the genetic component of resistance variation, we used a set of 12 inbred lines. The variation among line means provides an estimate of the standing genetic variation segregating with the original population and the variation among individuals within lines is an estimate of the environmental or experimental variation. Each inbred line was exposed, in replicate, to the population-by-insecticide specific LD_50_ to determine whether genetic variation was segregating in the population for susceptibility/resistance to each insecticide.

We detected significant heritable genetic variation in all 25 insecticide-by-population combinations with one exception, Tanzania treated with Sevin (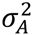 = 0.0027, SE +/− 0.0019, *p* = 0.1857; *h*^2^ = 0.4070, SE +/− 0.2514, *p* = 0.100). Despite having the longest history of past selection and some of the highest values of LD_50_, Malathion heritabilities were extremely high ( > 0.9) in Purdue (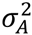 = 0.0749, SE +/− 0.0350, *p* = 0.0554; *h*^2^ = 0.9790, SE +/− 0.0187, *p* << 0.0001), Tanzania (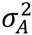 = 0.1012, SE +/− 0.0208, *p*< 0.0005; *h*^2^ = 0.9625, SE +/− 0.0268, *p* << 0.0001), and c-SM (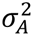 = 0.1024, SE +/− 0.0294, *p* = 0.0025; *h*^2^ =0.9633, SE +/− 0.0426, *p* << 0.0001). The insecticide with the largest range of heritabilities was the recently introduced Imidacloprid with Bhopal having the highest heritability (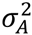 = 0.0201, SE +/− 0.0096, *p* = 0.0596; *h*^2^ = 0.8113, SE +/− 0.0978, *p* << 0.0000) and Jerez (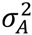 = 0.0014, SE +/− 0.0006, *p* = 0.0396; *h*^2^ = 0.2927, SE +/− 0.0708, *p* = 0.0017) the lowest. We found that *h*^2^ was not correlated with a population's mean resistance, measured by its LD_50_ (*r*[LD_50_, *h*^2^] = 0.0405, n = 25, N.S.). Thus, no matter how high the mean resistance of a population to an insecticide, genetic variation for further resistance remained within it.

### Genetic correlations among resistance phenotypes

Estimation of a genetic correlation between two traits generally requires measuring the two phenotypes on the same individual. This is not possible when the phenotypes are deaths on two insecticides. However, because individuals from the same inbred line are nearly genetically identical, we measured the response to two separate insecticides on different individuals from the same set of inbred lines, whose degree of genetic relatedness is known. In this way, the correlation between inbred line means can be related quantitatively to the genetic variation and covariation of individual traits in the population from which the inbred lines were derived. Using this method, we estimated the genetic correlations reported in Tables 3 and 4.

**Table 3.**
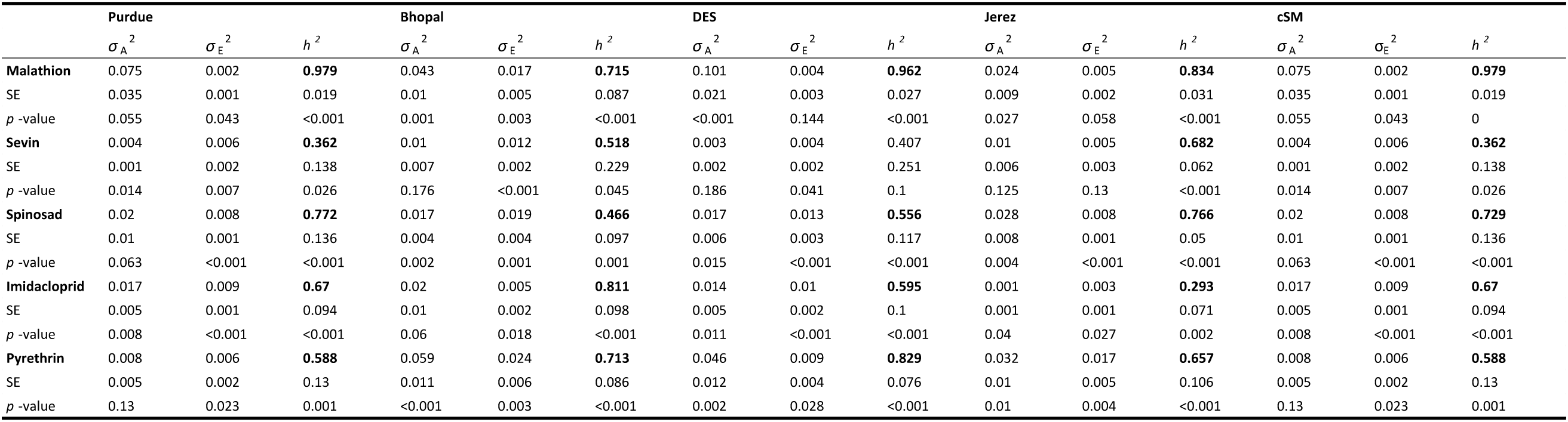
The additive variance (σ_A_^2^), the environmental or error variance (σ_E_^2^) and the heritability (*h*^2^) of the resistance phenotype for each insecticide applied to each population at the LD50 of that population.

**Table 4.** The additive variance (σA2), the environmental or error variance (σE2) and the heritability (h2) of the resistance phenotype for each insecticide applied to each population at the LD90 of that population.

|  | Purdue |  |  | Bhopal |  |  |
| --- | --- | --- | --- | --- | --- | --- |
| | $\sigma A^2$ | $\sigma E^2$ | $h^2$ | $\sigma A^2$ | $\sigma E^2$ | $h^2$ |
| **Imidacloprid** | 0.015 | 0.014 | **0.522** | 0.002 | 0.003 | **0.391** |
| SE | 0.007 | 0.004 | 0.144 | 0.001 | 0.001 | 0.103 |
| *p*-value | 0.027 | 0.005 | 0.001 | 0.009 | 0.002 | <0.001 |
| **Malathion** | 0.072 | 0.015 | **0.824** | 0.033 | 0.012 | **0.73** |
| SE | 0.03 | 0.005 | 0.053 | 0.008 | 0.004 | 0.103 |
| *p*-value | 0.025 | 0.022 | <0.001 | 0.002 | 0.012 | <0.001 |
| **Sevin** | 0.003 | 0.006 | *0.341* | 0.008 | 0.011 | **0.402** |
| SE | 0.002 | 0.002 | 0.154 | 0.004 | 0.003 | 0.118 |
| *p*-value | 0.098 | 0.005 | 0.055 | 0.095 | 0.005 | <0.001 |
| **Spinosad** | 0.018 | 0.011 | **0.625** | 0.015 | 0.019 | **0.443** |
| SE | 0.011 | 0.003 | 0.195 | 0.004 | 0.004 | 0.11 |
| *p*-value | 0.144 | 0.003 | 0.004 | 0.002 | 0.001 | <0.001 |

Genetic correlations estimated near the parent population LD_50_ ranged from a high of +0.840 to Sevin and Imidacloprid sensitivity for the Jerez population to a low of -0.628 for the Purdue population between sensitivity to Sevin and Malathion (Table 3). Genetic correlations between the same two insecticides varied widely among populations. In the most extreme case, resistance to Malathion and Sevin were highly positively correlated (+0.796) in the Tanzania population but highly negatively correlated (-0.620) in the Jerez population. Thirty-eight percent of the 50 correlations estimated were significant. Among the 19 significant genetic correlations, positive genetic correlations outnumbered negative correlations 16 to 3.

### Stability of correlations among populations

A random skewers test (see above) showed that the genetic variance-covariance matrixes for the Purdue, Bhopal and Tanzania populations were significantly similar to one another (Table 6). Jerez was also quantitatively similar to these but not significantly so, while the genetics of the laboratory c-SM population were the least similar to those of the four wild populations.

### Stability of correlations among concentrations

Only two populations, Purdue and Bhopal, were investigated for genetic correlations in the LD_90_ response of inbred lines to multiple insecticides (Table 4). The correlation between inbred line mean responses to an insecticide at its LD_50_ and LD_90_ tended to be high and positive (Table 5): five of the eight measured correlations were significantly positive, while another was borderline significant (*p* = 0.052). A random skewers test (see above) showed that genetic variance-covariance matrixes for the Purdue population estimated at the LD_50_ and LD_90_ were essentially identical (*r* = +0.95, *p* = 0,008). Those of Bhopal were similar but not significantly so (Table 6).

**Table 5.**
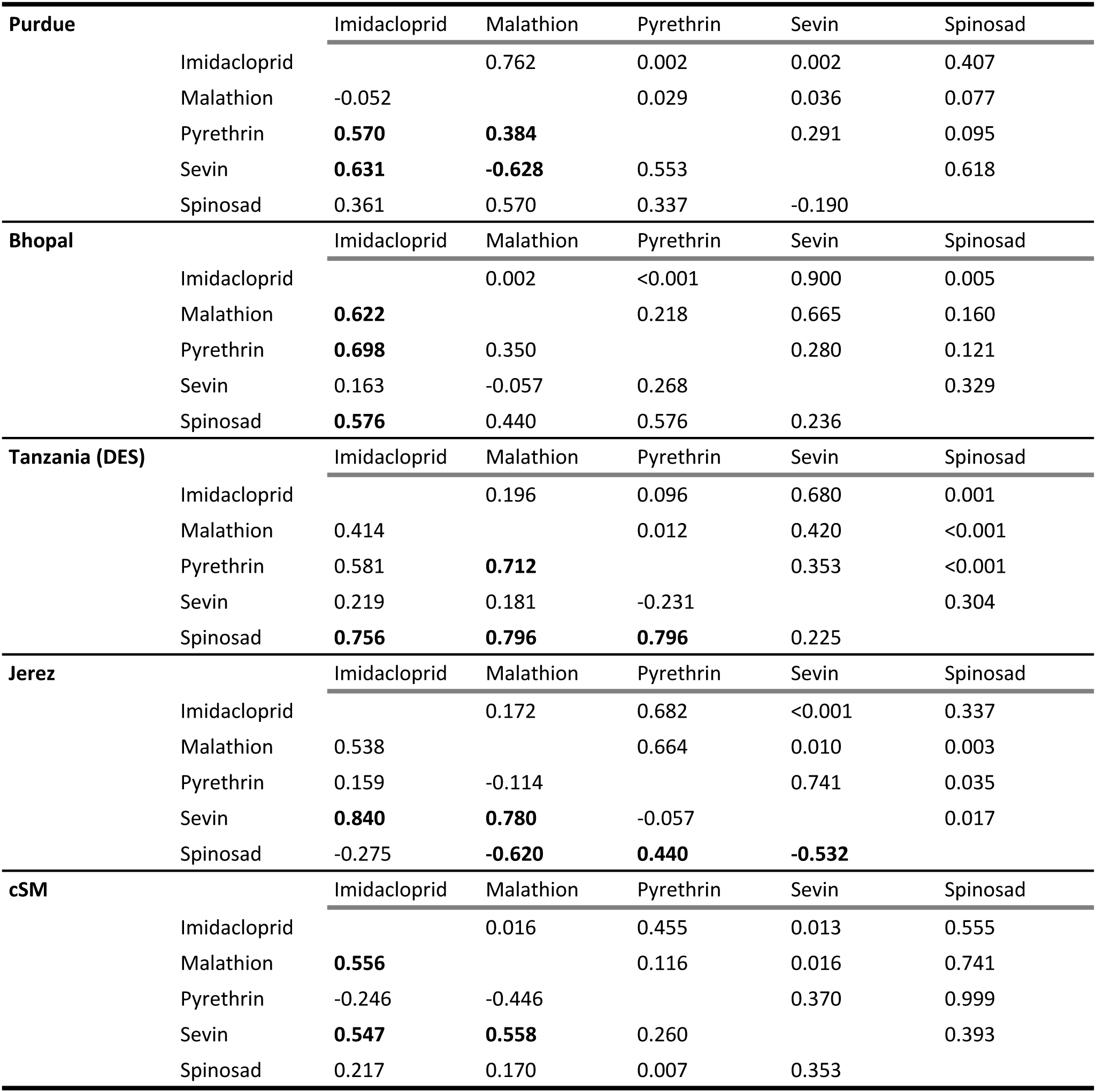
Genetic correlations (*rG*) between insecticides for each population at the LD_50_ below the diagonal and the significance level (*p*) above the diagonal.

**Table 6.** Genetic correlations (*rG*) between insecticides for each population at the LD90 below the diagonal and the significance level (*p*) above the diagonal. (No values for Pyrethrin are included due to timing constraints)

| **Purdue** |  | **Imidacloprid** | **Malathion** | **Sevin** | **Spinosad** |
| --- | --- | --- | --- | --- | --- |
|  | **Imidacloprid** |  | 0.801 | 0.082 | 0.839 |
|  | **Malathion** | 0.121 |  | 0.749 | 0.065 |
|  | **Sevin** | 0.480 | -0.121 |  | 0.590 |
|  | **Spinosad** | -0.090 | 0.676 | -0.182 |  |
| **Bhopal** |  | **Imidacloprid** | **Malathion** | **Sevin** | **Spinosad** |
|  | **Imidacloprid** |  | 0.263 | 0.502 | 0.386 |
|  | **Malathion** | -0.313 |  | 0.021 | <0.001 |
|  | **Sevin** | 0.229 | **0.480** |  | 0.708 |
|  | **Spinosad** | 0.308 | **-0.835** | -0.103 |  |

### Projection and predictive intervals

We used the estimated narrow sense heritabilities, *h^2^* (Tables 1 and 2), and the genetic correlations (Tables 3 and 4) to project the evolution of insecticide resistance in response to the application of one insecticide or a rotation between two pesticides at the population’s LD_50_. We report evolutionary change due to insecticide treatment as the fractional change in the LD_50_ in Figure 3. We used the genetic correlations with heritabilities to estimate the expected change in LD_50_ of one insecticide due to the treatment of the population with a different insecticide. As resistance evolves, application of an LD_50_ insecticide concentration becomes increasingly futile (Table 9). From the stochastic projections, using parameters estimated from the Purdue population, Malathion resistance would more than double (1.011-fold increase, 95% CI: 0.83-1.14) in just 5 generations of exposure at its LD_50_. Similarly, the two most recently approved insecticides would have a greater that 50% increase in their resistance profiles after 5 generations of treatment (Imidacloprid: 0.563-fold increase, 95% CI: 0.32-0.82; Spinosad: 0.691-fold increase, 95% CI: 0.32-1.07). We also estimated *h*^2^ and *r_G_* for two populations at the LD_90_ concentration. We find that heritabilities and genetic correlations between treatments to remain constant across concentrations (Table 5) as well as between populations (Table 6).

**Figure 2.**
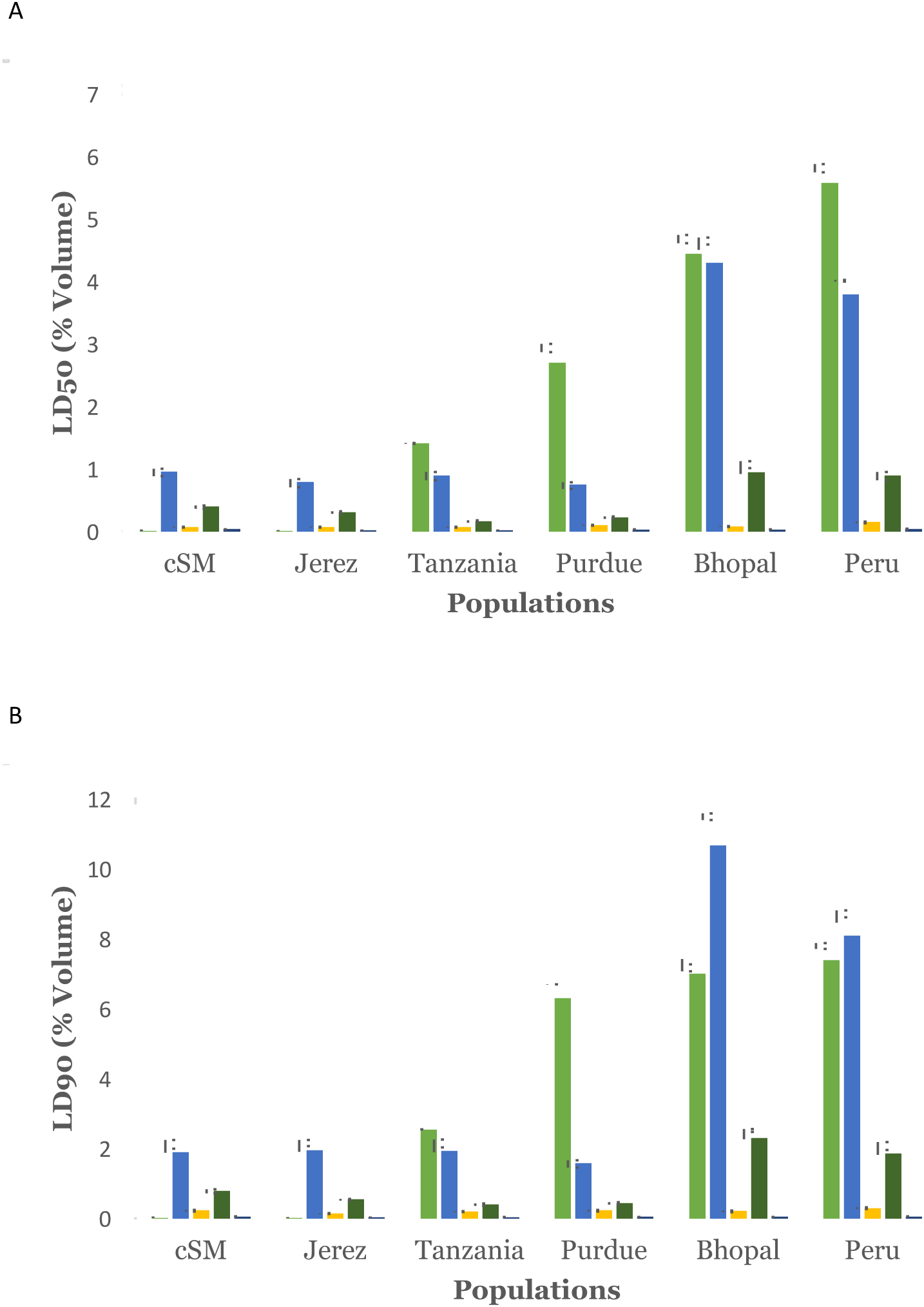
The average concentrations in percent by volume of the insecticides, reading from left to right Malathion (light green), Sevin (blue), Spinosad (yellow), Imidacloprid (dark green), Pyrethrin (dark blue), required to kill 50% (LD_50_, A) or 90% (LD_90_, B) of adult beetles from the six populations listed along the x-axis.

**Figure 3.**
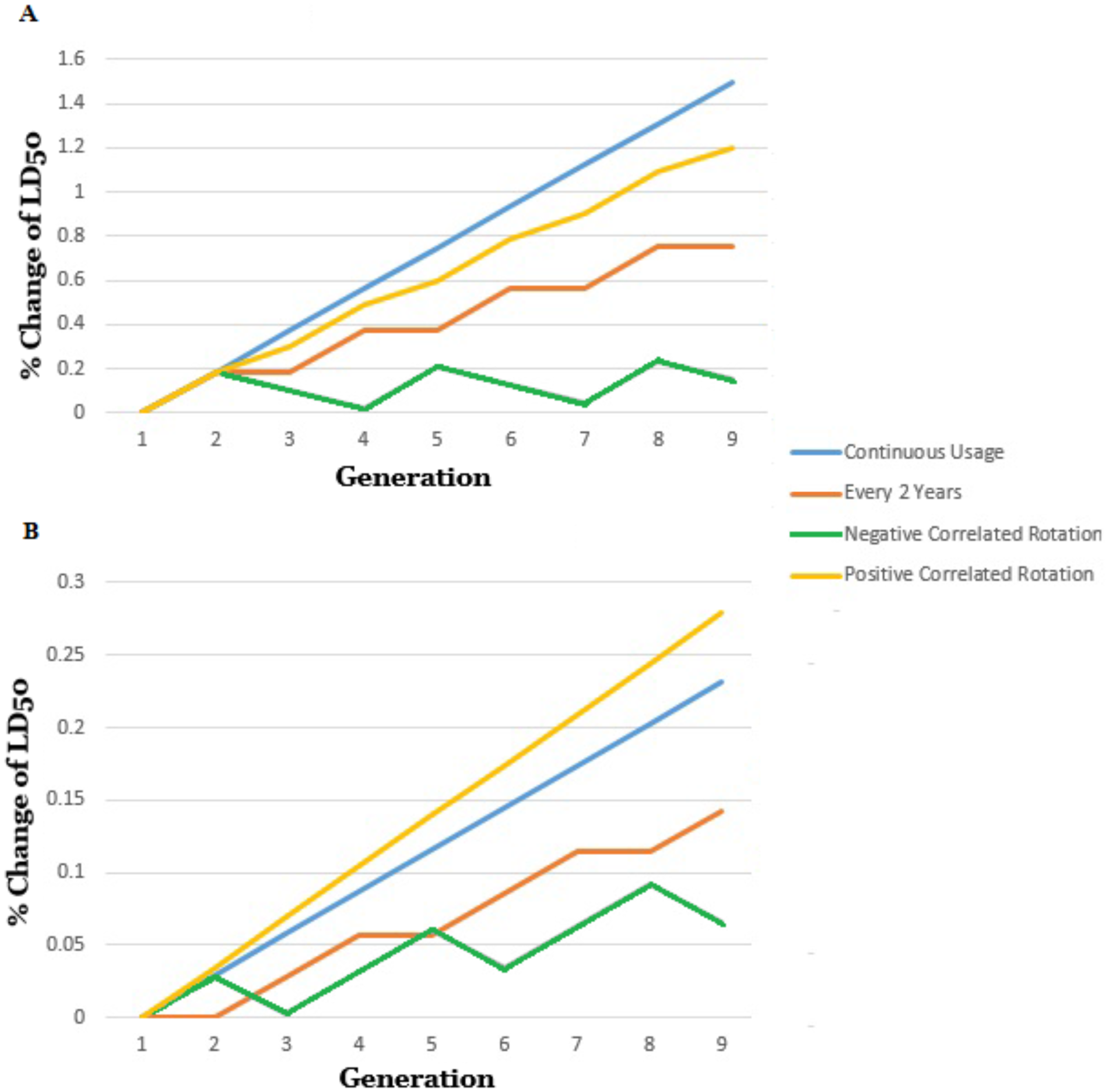
The estimated effects of four separate insecticide rotation strategies on the evolution of resistance towards Malathion (A) and Sevin (B). 1) Continuous spraying with no rotation (blue). 2) Spraying one insecticide every two years with no rotation (orange). 3) Yearly rotation between negatively correlated insecticides (gray). 4) Yearly rotation between positively correlated insecticides.

**Table 7.** Correlation between response at LD50 and response at LD90. (No values for Pyrethrin are included because at the higher dosage it was lethal to all beetles from all populations)

|  | **Purdue** |  | **Bhopal** |  |
| --- | --- | --- | --- | --- |
|  | Covariance | Correlation | Covariance | Correlation |
| **Imidacloprid** | 0.008 | **0.510** | 0.004 | 0.571 |
| SE | 0.004 | 0.213 | 0.002 | 0.234 |
| $p$-value | 0.093 | 0.043 | 0.843 | 0.052 |
| **Malathion** | 0.052 | **0.697** | -0.016 | -0.428 |
| SE | 0.023 | 0.167 | 0.009 | 0.245 |
| $p$-value | 0.060 | 0.002 | 0.137 | 0.165 |
| **Sevin** | 0.002 | **0.606** | 0.007 | **0.852** |
| SE | 0.001 | 0.279 | 0.005 | 0.291 |
| $p$-value | 0.267 | 0.048 | 0.048 | 0.004 |
| **Spinosad** | 0.002 | 0.114 | 0.013 | **0.900** |
| SE | 0.006 | 0.319 | 0.004 | 0.051 |
| $p$-value | 0.724 | 0.863 | 0.205 | <0.001 |

**Table 8.** Similarity among genetic varience-covariance matrices with respect to their predicted evolutionary responses estimated by the method of random skewers.

| **G-matrix Similarity Among Populations** |  |  |  |  |  |
| --- | --- | --- | --- | --- | --- |
|  | Purdue | Bhopal | Des | Jerez | cSM |
| Purdue |  | 0.064 | 0.031 | 0.214 | 0.182 |
| Bhopal | **0.695** |  | 0.026 | 0.081 | 0.212 |
| Des | **0.795** | **0.815** |  | 0.125 | 0.224 |
| Jerez | 0.397 | 0.657 | 0.55 |  | 0.179 |
| cSM | 0.451 | 0.409 | 0.372 | 0.456 |  |

| **G-matrix Consistency Among Lethal Doses (50:90)** |  |  |
| --- | --- | --- |
|  | r | p |
| Purdue | **0.95** | 0.008 |
| Bhopal | 0.53 | 0.17 |

**Table 9.** Means and 95% predictive intervals for stocastic simulations of the fractional increase of insecticide resistence when the Purdue populations is exposed to the chemical at its median lethal dose.

|  | **Generation of Exposure** |  |  |  |  |
| --- | --- | --- | --- | --- | --- |
|  | **1st** | **2nd** | **3rd** | **4th** | **5th** |
| **Imidacloprid** | 0.112 | 0.225 | 0.338 | 0.450 | 0.563 |
| 2.50% | 0.01 | 0.08 | 0.15 | 0.24 | 0.32 |
| 97.50% | 0.23 | 0.39 | 0.54 | 0.68 | 0.82 |
| **Malathion** | 0.202 | 0.404 | 0.606 | 0.808 | 1.011 |
| 2.50% | 0.10 | 0.28 | 0.46 | 0.65 | 0.83 |
| 97.50% | 0.25 | 0.48 | 0.70 | 0.92 | 1.14 |
| **Pyrethrin** | 0.090 | 0.179 | 0.268 | 0.358 | 0.448 |
| 2.50% | -0.04 | -0.01 | 0.03 | 0.08 | 0.14 |
| 97.50% | 0.24 | 0.39 | 0.52 | 0.65 | 0.77 |
| **Sevin** | 0.046 | 0.092 | 0.138 | 0.184 | 0.230 |
| 2.50% | -0.07 | -0.07 | -0.06 | -0.05 | -0.03 |
| 97.50% | 0.18 | 0.27 | 0.35 | 0.43 | 0.51 |
| **Spinosad** | 0.138 | 0.276 | 0.415 | 0.553 | 0.691 |
| 2.50% | -0.02 | 0.05 | 0.13 | 0.23 | 0.32 |
| 97.50% | 0.32 | 0.52 | 0.71 | 0.90 | 1.07 |

**Table 10.** Means and 95% predictive intervals for stocastic simulations of the fractional increase of insecticide resistence when the Purdue populations is exposed to differing chemicals with know genetic correlations at its median lethal dose. Negative Correlation: Malathion-Seven, Positive Correlation: ImidaclopridSevin, No Correlation, Malathion-Spinosad.

|  | Generation of Exposure |  |  |  |  |
| --- | --- | --- | --- | --- | --- |
|  | 1st | 2nd | 3rd | 4th | 5th |
|  | Malathion | Sevin | Malathion | Sevin | Malathion |
| **Malathion** | 0.112 | 0.092 | 0.120 | 0.066 | 0.128 |
| 2.50% | 0.01 | -0.12 | -0.12 | -0.21 | -0.19 |
| 97.50% | 0.23 | 0.24 | -0.12 | 0.35 | 0.46 |
| **Sevin** | -0.054 | -0.011 | -0.065 | -0.022 | -0.076 |
| 2.50% | -0.20 | -0.19 | -0.30 | -0.28 | -0.37 |
| 97.50% | 0.09 | 0.18 | 0.17 | 0.24 | 0.22 |
|  | Imidacloprid | Sevin | Imidacloprid | Sevin | Imidacloprid |
| **Imidacloprid** | 0.113 | 0.167 | 0.228 | 0.282 | 0.344 |
| 2.50% | 0.01 | 0.03 | 0.02 | 0.06 | 0.07 |
| 97.50% | 0.23 | 0.31 | 0.45 | 0.52 | 0.63 |
| **Sevin** | 0.054 | 0.097 | 0.151 | 0.194 | 0.248 |
| 2.50% | -0.03 | -0.04 | -0.01 | 0.00 | 0.03 |
| 97.50% | 0.14 | 0.24 | 0.32 | 0.40 | 0.47 |
|  | Malathion | Spinosad | Malathion | Spinosad | Malathion |
| **Malathion** | 0.202 | 0.310 | 0.481 | 0.588 | 0.760 |
| 2.50% | 0.10 | -0.07 | 0.05 | 0.01 | 0.15 |
| 97.50% | 0.25 | 0.69 | 0.94 | 1.17 | 1.41 |
| **Spinosad** | 0.108 | 0.246 | 0.353 | 0.491 | 0.599 |
| 2.50% | -0.27 | -0.16 | -0.20 | -0.09 | -0.09 |
| 97.50% | 0.48 | 0.66 | 0.91 | 1.07 | 1.28 |

To illustrate how the efficacy of windowing depends jointly on the heritability (*h*^2^) and the genetic correlation (*r_G_*), we explored 4 different insecticide application strategies for controlling the Purdue population (see Figure 3). Resistance evolves most rapidly when an insecticide of high heritability is applied continuously, generation after generation (Table 9). When the same insecticide is applied periodically, resistance evolves at a fraction of the continuous rate. However, the evolution of resistance is nearly halted by windowing or rotating every generation between two insecticides with a negative genetic correlation of similar magnitude to the heritability. For example, in simulations, rotating the insecticides Sevin and Malathion results in significantly impeding the evolution of resistance to both chemicals. By generation 5, the population has been exposed to Malathion 3 times and resistance to the chemical has only increased 12.8% (95% CI: -0.19-0.46). Whereas, if the population was treated with Malathion in 3 sequential generations the resistance profile would increase 60.6% (95% CI 0.46-0.70). A slowing of the evolution of resistance by 47.5%. The population would have been exposed to Sevin twice, and because of the negative correlation between the two insecticides, the population actually becomes more susceptible to Sevin treatment ( -0.076-fold change, 95% CI -0.37-0.22).

However, windowing between insecticides with a positive genetic correlation is comparable to continually applying a single insecticide. In the case where the genetic correlation is positive and larger than the heritability, windowing could accelerate the rate at which resistance evolves. For example, rotating Imidacloprid with Sevin actually hastens the evolution of resistance to Sevin. After five generations of rotation (3 Imidacloprid, 2 Sevin) there is a 0.248-fold increase (95% CI 0.03-0.47) in resistance to Sevin when just 2 sequential treatments of Sevin by itself only produces a 0.092-fold increase (95% CI -0.07-0.27). This change in resistance to Sevin is higher that of 5 sequential Sevin treatments alone (0.230-fold increase, 95% CI-0.03-0.51).

## Discussion

Current agricultural practices rely heavily on insecticides to control most pest species. As every economically significant insect order evolves resistance to these chemicals, it is imperative to develop novel genetic tools for pest control to preserve crop yields and limit disease transmission. Newer strategies of pest control, such as transgenic crops and endonuclease and microorganism mediated gene drives (Esvelt et al. 2014), aim to decrease dependence on insecticides, which are becoming less effective and can have negative side effects on particular ecological communities (Whitehorn et al. 2012). While these technologies are in the process of development, insecticide-usage strategies like windowing may prolong the efficacy of the current pest control methods, allowing for more time for innovation while still maintaining economic yields.

We find that populations are highly variable from one another in their sensitivity to insecticides and the data suggest that some of this variation is an evolutionary product of their history of exposure to insecticides. Specifically, two of our wild populations, Bhopal and Peru, have had a history of a brief but intense exposure (Bhopal) or long term exposure (Peru) to Malathion and chemically related insecticides. Beetles from both populations are much more resistant to pesticides overall than are beetles from c-SM, the naïve laboratory population. Another aspect of our findings also indicates the importance of a population’s history of exposure to insecticides. In general, among the five wild populations, we observed significantly greater resistance to the older insecticides, Malathion and Sevin, and much lower resistance to the more recently introduced insecticides, Imidacloprid and Spinosad. Moreover, the variation among populations in resistance is greater for the older insecticides and much lower for those more recently introduced.

The large differences in resistance between and within populations allowed us to estimate the heritability of resistance to each insecticide as well as the genetic correlation of resistance across insecticides. These genetic estimates allowed us to test the efficacy of the pest control strategy of ‘windowing,’ whose efficacy depends upon rotating the application of negatively genetically correlated insecticides. Surprisingly, we found instances of negative genetic correlations in resistance to insecticides sharing the same mechanism of action (Purdue population) as well as the converse, namely, positively correlated insecticides acting on entirely different pathways (Spinosad and Pyrethrin resistance in the Tanzania population). Although integrated pest management strategies have been designed to insure rotation of insecticides with different modes of action, the evolution of insecticide resistance depends critically on the sign and magnitude of the genetic correlation of resistance between the rotated insecticides. As we showed in Figure 7, a negative genetic correlation is critical to limiting or preventing the evolution of insecticide resistance. Rotation between insecticides with a positive genetic correlation of pest resistance can lead to more rapid evolution of resistance to both pesticides than would the application of only a single pesticide. Unfortunately, our data indicate that the sign and magnitude of a genetic correlation depends not only on the pair of insecticides to be rotated but also on the specific host population targeted for control. That is, rotating between Malathion and Sevin would limit the evolution of resistance in the Purdue population, where the genetic correlation of resistance is negative, but rotation of the same two insecticides would accelerate the evolution of resistance in the Tanzania population, where the genetic correlation is positive.

We estimated heritability and genetic correlations at both the LD_90_ as well as the LD_50_ concentration for all insecticides for two pest populations and found that the genetic correlations of resistance were nearly constant across the two concentrations. This finding is important, if it proves to be general. Constancy of the genetic correlations of resistance across insecticide concentrations permits the development of a management strategy that remains effective even as resistance profiles evolve. Our findings support insecticide rotation as an efficacious strategy for limiting or halting the evolution of insecticide resistance when the genetic correlations among the rotated insecticides are negative. Therefore, we recommend the estimation of the heritability and genetic correlations of resistance between insecticides and to develop application strategies based on negatively correlated genetic responses while avoiding simultaneous or sequential use of insecticides for which pest resistance is positively genetically correlated.

## Acknowledgments

This work was supported by Indiana University and the National Institutes of Health R01 GM084238 (M.J.W.).

---

**Supp. Table 1.** Covariation of each insecticide pair for each population at the LD50. Below the diagnal: Estimate of the additive genetic covarience between the traits; Above the daignal: *p* value of one sample, two tailed t-test for difference from 0.

| Purdue |  | Imidacloprid | Malathion | Pyrethrin | Sevin | Spinosad |
| --- | --- | --- | --- | --- | --- | --- |
|  | Imidacloprid |  | 0.848 | 0.079 | 0.023 | 0.378 |
|  | Malathion | -0.002 |  | 0.118 | 0.139 | 0.168 |
|  | Pyrethrin | 0.007 | 0.011 |  | 0.436 | 0.202 |
|  | Sevin | 0.005 | -0.010 | 0.002 |  | 0.661 |
|  | Spinosad | 0.007 | 0.022 | 0.004 | -0.001 |  |
| Bhopal |  | Imidacloprid | Malathion | Pyrethrin | Sevin | Spinosad |
|  | Imidacloprid |  | 0.033 | 0.030 | 0.677 | 0.047 |
|  | Malathion | 0.018 |  | 0.253 | 0.859 | 0.157 |
|  | Pyrethrin | 0.024 | 0.017 |  | 0.422 | 0.101 |
|  | Sevin | 0.002 | -0.001 | 0.006 |  | 0.427 |
|  | Spinosad | 0.011 | 0.012 | 0.018 | 0.003 |  |
| Des |  | Imidacloprid | Malathion | Pyrethrin | Sevin | Spinosad |
|  | Imidacloprid |  | 0.223 | 0.122 | 0.334 | 0.084 |
|  | Malathion | 0.016 |  | 0.030 | 0.458 | 0.024 |
|  | Pyrethrin | 0.015 | 0.048 |  | 0.444 | 0.024 |
|  | Sevin | 0.001 | 0.003 | -0.003 |  | 0.327 |
|  | Spinosad | 0.011 | 0.022 | 0.022 | 0.002 |  |
| Jerez |  | Imidacloprid | Malathion | Pyrethrin | Sevin | Spinosad |
|  | Imidacloprid |  | 0.274 | 0.591 | 0.151 | 0.446 |
|  | Malathion | 0.003 |  | 0.628 | 0.135 | 0.066 |
|  | Pyrethrin | 0.001 | -0.003 |  | 0.783 | 0.080 |
|  | Sevin | 0.003 | 0.012 | -0.001 |  | 0.168 |
|  | Spinosad | -0.002 | -0.016 | 0.013 | -0.009 |  |
| cSM |  | Imidacloprid | Malathion | Pyrethrin | Sevin | Spinosad |
|  | Imidacloprid |  | 0.093 | 0.492 | 0.056 | 0.485 |
|  | Malathion | 0.023 |  | 0.089 | 0.084 | 0.662 |
|  | Pyrethrin | -0.007 | -0.033 |  | 0.420 | 0.994 |
|  | Sevin | 0.014 | 0.036 | 0.011 |  | 0.303 |
|  | Spinosad | 0.004 | 0.008 | 0.000 | 0.010 |  |

**Supp. Table 2.**
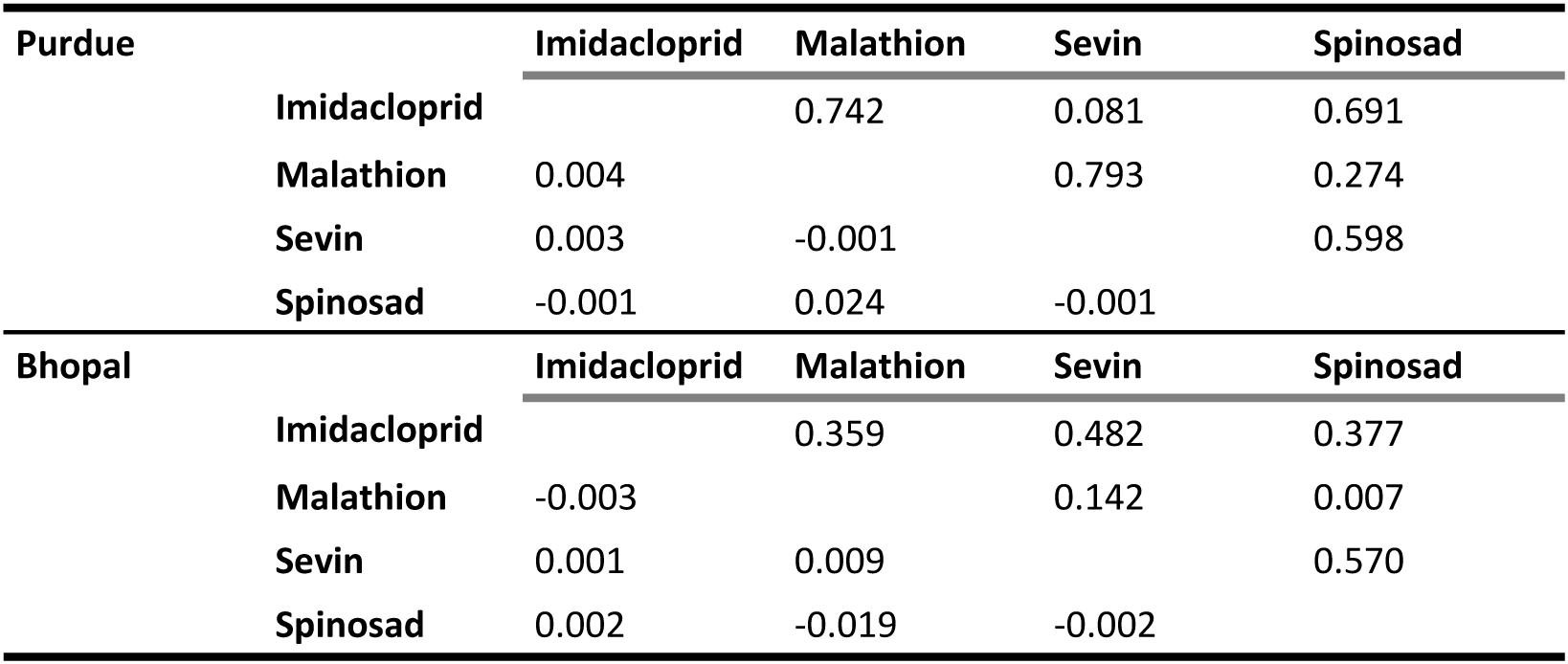
covariation of each insecticide pair for each population at the LD_90_. Below the diagnal: Estimate of the additive genetic covarience between the traits; Above the daignal: *p*-value of one sample, two tailed t-test for difference from 0.

| **Purdue** |  | **Imidacloprid** | **Malathion** | **Sevin** | **Spinosad** |
| --- | --- | --- | --- | --- | --- |
|  | **Imidacloprid** |  | 0.742 | 0.081 | 0.691 |
|  | **Malathion** | 0.004 |  | 0.793 | 0.274 |
|  | **Sevin** | 0.003 | -0.001 |  | 0.598 |
|  | **Spinosad** | -0.001 | 0.024 | -0.001 |  |
| **Bhopal** |  | **Imidacloprid** | **Malathion** | **Sevin** | **Spinosad** |
|  | **Imidacloprid** |  | 0.359 | 0.482 | 0.377 |
|  | **Malathion** | -0.003 |  | 0.142 | 0.007 |
|  | **Sevin** | 0.001 | 0.009 |  | 0.570 |
|  | **Spinosad** | 0.002 | -0.019 | -0.002 |  |

**Supp Figure 1.**
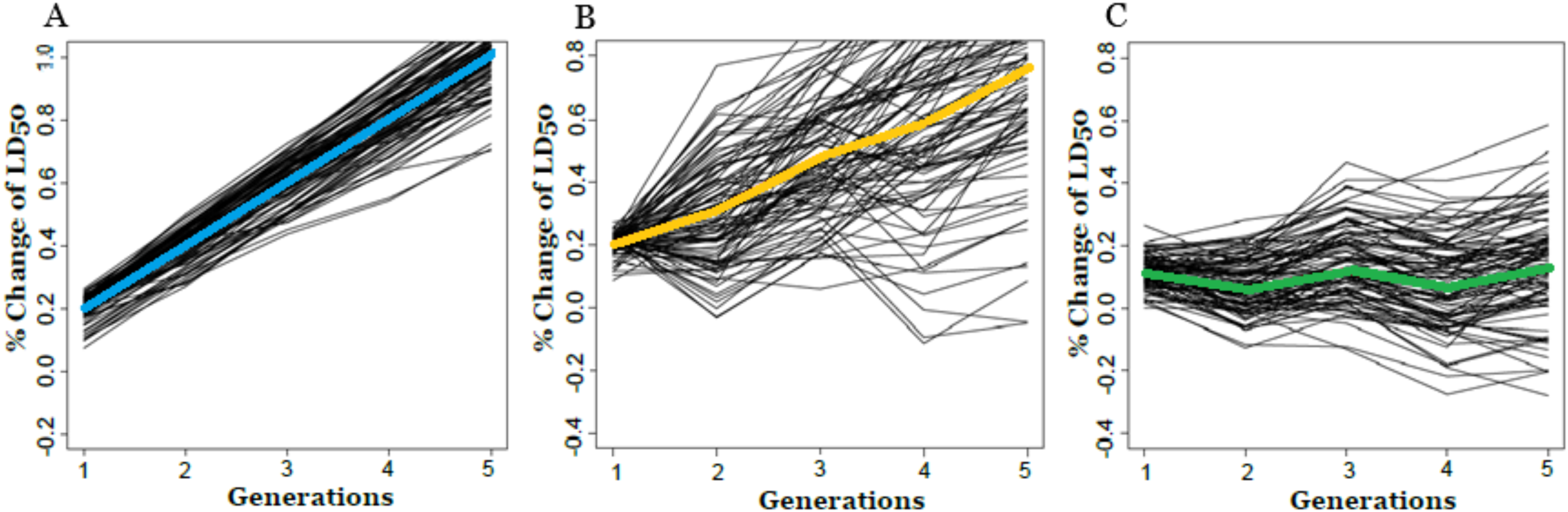
The variance about the estimated effects of four separate insecticide rotation strategies on the evolution of resistance towards Malathion (A) and Sevin (B). 1) Continuous application of Malathion with no rotation (blue). 2) application of one insecticide, Malathion, every other year with no rotation (orange). 3) Yearly rotation between negatively correlated insecticides, Malthion and Sevin (gray).

